## Supplemental Information for "Dual role of Miro protein clusters in mitochondrial cristae organisation and ER-Mitochondria Contact Sites"

<u>This file contains:</u>

**Supplementary Figure and Table legends**

**Supplementary Material and Methods**

**Supplementary References**

**Supplementary Table 1**

**Supplementary Figures**

**Supplementary Figures and Table legends**

**Supplementary Table 1:** Complete antibody information including provider, clone and/or catalogue number and dilution used in each figure panel.

**Supplementary Fig. 1:** Super-resolution imaging of mitochondrial matrix. Rescue of matrix morphology upon overexpression of either of Miro proteins. DKO cells were transfected with $^{Myc}$Miro1 or $^{Myc}$Miro2 along with mtRo-GFP and imaged under a structured illumination microscope. Scale bar: 5 μm.

**Supplementary Fig. 2:** HeLa cells were transfected with mitochondrially targeted $^{GFP}$Su9 or $^{GFP}$Miro1 and immunoprecipitated using GFP trap beads. While $^{GFP}$Miro1 specifically pulls down MICOS components, $^{GFP}$Su9 did not show any interactions with MICOS proteins.

**Supplementary Fig. 3:** $^{Myc}$Miro1 transfected HeLa cells were immunostained with anti-Myc antibody followed by Alexa 488 conjugated secondary antibody and imaged using a Structured Illumination Microscope using 488 nm laser. Z stacks were collected and merged using maximum intensity projection. Scale bar: 10 μm, Inset: 3 μm.

**Supplementary Fig. 4:** SIM imaging of endogenous Miro2 in HeLa cells. Tom70(1-70)$^{GFP}$ transfected cells were labelled with Miro2 antibody followed by secondary antibodies conjugated to Alexa 555. Z stacks were acquired and merged using maximum intensity projection in ImageJ. Scale bar. 1 μm.

**Supplementary Fig. 5:** Correlated SIM and dSTORM imaging of $^{Myc}$Miro1. HeLa cells were stained with anti-Myc antibody followed by simultaneous labelling with Alexa 488 and Alexa 647 conjugated secondary antibodies. SIM was performed with 488 nm laser while dSTORM of the same cell was performed using 642 nm laser. Inset: Magnification of the boxed region. Scale Bar: 20 μm, Inset: 5 μm.

**Supplementary Fig. 6:** dSTORM imaging of HeLa cells differently expressing $^{GFP}$Miro2. Three different expression level: low, moderate and high, were chosen to demonstrate cluster formation by $^{GFP}$Miro2. Scale bar: 5 μm (1 μm in zoom images).

**Supplementary Fig. 7**: (A) MEFs were transfected with $^{GFP}$Miro2, immunostained with anti-GFP antibody followed by Alexa 647 conjugated secondary antibody and imaged using a custom built TIRF based microscope with 642 nm laser. (B) dSTORM image of mitochondria in a rat hippocampal neuron transfected with $^{GFP}$Miro2. Enlarged image of the boxed region shown in (A, B). Scale bar: 5 μm, Inset: 0.5 μm.

**Supplementary Fig. 8:** (A, B) Immunofluorescence of HeLa cells transfected with $^{GFP}$Miro1 (A) or $^{GFP}$Miro2 (B) with MINOS complex protein Mitofilin. Cells were stained with anti-GFP and anti-Mitofilin antibodies and imaged using a confocal microscope. (C) Nearest-Neighbour distances between Miro2 and Mic60/Mitofilin. Nearest-Neighbour distances were calculated from the reconstructed images of Miro and Mic60/Mitofilin by detecting each cluster with Mosaic Interaction analysis plugin available with ImageJ. Scale bar: 10 μm.

### Supplementary Materials and Methods

#### Cell culture and transfection

HeLa and HEK293T cells were cultured in DMEM medium (Gibco) supplemented with streptomycin (100 µg/ml), penicillin (100 U/ml), and 10% fetal bovine serum in a 10 cm cell culture dishes. Mouse Embryonic Fibroblast cells were cultured similarly in 15% fetal bovine serum. Cells were transfected with ~5 µg plasmid DNA using nucleofection (Amaxa, Lonza AG) according to manufacturer's protocol. Post-transfection, cells were plated onto 25 nm PLL coated (HeLa and HEK293T cells) or 20µg/ml Fibronectin coated coverslips (Sigma) (MEF cells) containing with 100 nm TetraSpeck™ fluorescent Microspheres (Life Technologies).
Rat hippocampal cultures were performed as described earlier (Macaskill, Rinholm et al., 2009). Neurons were transfected at 7-9 DIV using Lipofectamine®2000 in complete media. Typically, 3 µl Liopfectamine was complexed with 5 µg plasmid DNA in neurobasal media containing 0.59% D-Glucose. 30 min post-complexation DNA and Lipofectamine mixture was diluted in complete culture media and added to the wells of a 6 well plate containing neurons. Media was replaced after 2 h and further cultured for 2-3 days before fixation.

#### Plasmid DNA, antibodies and reagents

$^{Myc}$Miro1 and $^{Myc}$Miro2 plasmids were obtained from Prof. Pontus Aspenström (Karolinska Institutet)(Fransson, Ruusala et al., 2003). $^{GFP}$Miro1 and $^{GFP}$Miro2 were generated as described earlier (Birsa, Norkett et al., 2014). $^{Myc}$Mfn1 (plasmid #23212), $^{Myc}$Miro2ΔTM (Plasmid #47901), $^{GFP}$Su9 (Plasmid #23214) were obtained from Addgene (Chen, Detmer et al., 2003, Fransson, Ruusala et al., 2006). Tom70(1-70)$^{GFP}$ was described before (Covill-Cooke, 2017). EGFP-C1 plasmid was purchased from clonetech. KDEL-HRP construct was a gift from Dan Cutler's lab (Connolly, Futter et al., 1994). Anti-Miro1 (Atlas antibodies recognizing Miro1 and Miro2, 1 in 1000, Rabbit), anti-Miro2 (Neuromab, clone N384/63, 1 in 1000 for immunoblotting and 1 in 100 for immunofluorescence, Mouse), anti Myc (Neuromab clone 9E10, 1 in 100 for immunoblotting, 1 in 50 for immunofluorescence, Mouse), anti-cMyc (Santa Cruz Biotechnology, 1 in 50 for immunofluorescence, Rabbit), anti-Mitofilin (Abcam, 1 in 1000 for immunoblotting, 1 in 300-500 immunofluorescence, Rabbit), anti-CHCHD3 (Atlas antibodies 1 in 1000 for immunoblotting, 1 in 100-200 for immunofluorescence, Rabbit), anti-Sam50 (Atlas antibodies, 1 in 100 for immunoblotting Rabbit), anti-Sam50 (Abcam, 1 in 500 for immunoblotting, rabbit), Anti Metaxin1 (Atlas antibodies, 1 in 200 for immunoblotting, 1 in 30 for immunofluorescence, Rabbit), anti-Tom20 (Santacruz biotechnology, 1 in 500 for immunofluorescence), anti-GFP (Neuromab clone N86_38, 1 in 100 (Supernatant) for immunoblotting and 1 in 100 (Purified) for immunofluorescence, Mouse), anti-GFP nanobody (Chromotek GmBH). All secondary antibodies (Alexa 488, Alexa 555 and Alexa 647 conjugated, Life technologies) were used in 1 in 500 dilution unless otherwise specified. Alexa 647 conjugated anti-GFP nanobody was prepared using Microscale Protein Labeling Kit (Life technologies) according to manufacturer's protocol and purified using 3kDa molecular weight cut-off filters (Sigma).

**Immunostaining**

24 h post transfection, cells were fixed with 4% PFA (4% paraformaldehyde, 4% sucrose in 1× PBS, pH 7.0) prewarmed at 37$^{o}$C for 10 min. After four washes in 2 ml 1× PBS, coverslips were incubated for 30-45 min in blocking solution (1% BSA, 10% horse serum, 0.2% Triton X-100 in 1× PBS). Coverslips were then incubated for 1½ h with primary antibodies diluted in blocking solution, washed 8 times (each dipping 10 times) in 1× PBS, and incubated with secondary antibodies. After a further 10 washes (each dipping 10 times) in 1× PBS, coverslips were transferred in a 10 cm dish containing 15 ml 1× PBS and washed twice for 10 min each with constant agitation. Coverslips were further fixed for 8 min in 4% PFA and washed four times in 1× PBS. Cells were mounted in Prolong Gold (for Confocal and SIM) antifade reagent or 10% Vectashield H-1000 (Vector labs) in 95% Glycerol +50 mM TRIS, pH 8.0 buffer for dSTORM and correlated SIM-dSTORM techniques) and sealed using nail varnish.

**Biochemical assays**

For co-immunoprecipitation experiments, 24 h post transfection cells were lysed in 50 mM HEPES, pH 7.4, 150 mM NaCl, 1 mM EDTA, 1% Triton X-100, 1 mm PMSF, antipain, leupeptin, and pepstatin at 10 μg/ml each. After clearing the lysates of nuclei and cellular debris by centrifugation at 14000 rpm for 30 min, the samples were incubated with GFP-TRAP beads (Chromotek GmBH) for 2 h at 4$^{o}$C with constant rotation; the immunoprecipitated complexes were washed three times with the above buffer, eluted with Laemmli sample buffer and run in an acrylamide gel. Approximately 2% of the cell lysate used for the immunoprecipitations was used to load the inputs. Samples were then western-blotted on nitrocellulose membrane (GE Healthcare), developed with Crescendo substrate (Millipore), and detected with a LAS 4000 imager (GE Healthcare). For BN-Page, HeLa and MEF cell lysate was prepared using native page sample buffer containing 1% Digitonin (Life technologies), loaded onto 3-12% Bis-Tris polyacrylamide gel (Novex, life technologies) and electrophoresis was performed according to manufacturer's protocol. After resolving the protein complexes, western-blotting was performed on PVDF membrane (GE healthcare), fixed with 10% acetic acid and developed as described earlier.

**PLA assay**

MEFs were seeded on fibronectin coated coverslips and grown overnight. Coverslips were fixed at room temperature for 7 minutes using 4% paraformaldehyde in PBS (pH 7.5) supplemented with 4% sucrose, washed 3 times in PBS, and blocked for 10 minutes in blocking buffer (10% Horse Serum, 5 mg/ml BSA, and 0.2% Triton in PBS). Coverslips were then incubated for 1h at room temperature with primary antibodies in blocking buffer (1:200 Mouse-anti-Mitofilin (Abcam, clone 2E4AD5) and 1:200 Rabbit-anti-Samm50 (Atlas, HPA034537); 1:200 Mouse-anti-Mitofilin and 1:200 Rabbit-anti-CHCHD3 (Atlas, HPA042935); or 1:150 Mouse-anti-Miro2 (Neuromab, N384/63) and 1:150 Rabbit-anti-CHCHD3 (Atlas, HPA042935); or 1:200 Rabbit-anti-Samm50 (Atlas, HPA034537)). The remaining of the staining was performed according to the manufacturer's protocol (Duolink® PLA). In brief, coverslips were incubated in a humidity chamber with anti-mouse MINUS and anti-rabbit PLUS probes (Sigma Aldrich) for 1h at 37ºC, hybridized for 30 minutes at 37ºC, and amplified for 100 mins at 37ºC using the red fluorophore. Coverslips were then allowed to dry at room temperature for 20 mins in the dark, and mounted using Duolink® In Situ mounting medium containing DAPI.

**Imaging**

**Confocal imaging**

Confocal imaging was performed in a Zeiss LSM 700 confocal microscope with a plan-apochromat 63× oil-immersion lens with 1.4 numerical aperture (NA). Z stacks were acquired with 0.5 µm step size and then merged using maximum intensity projection in ImageJ.

**SIM, correlated SIM and dSTORM, 3D dSTORM imaging**

Structured Illumination Microscopy was performed in Zeiss Elyra PS.1 equipped with 405, 488, 555 and 642 nm lasers. Images were acquired with 63×1.4 NA oil immersion objective using pco.edge sCMOS camera and Zen 2012 image analysis software. SIM imaging was performed with 34 µm grating with 3 rotations. Typically, images were acquired with 1-3%

laser intensity and 120-150 ms exposure time. SIM stacks were typically composed by 50 planes acquired at 0.14 μm steps corresponding to ~7-8 μm in depth.

Correlated SIM and dSTORM imaging was performed in the same microscope with 100×1.46 NA oil immersion objective. SIM images were acquired first with 488 laser line and then switched to dSTORM mode where images were acquired using 642 nm laser. Typically 15000 frames (50 ms per frame) in TIRF mode were acquired after initial bleaching and occasionally pulse of 405 nm laser was used to boost the blinking performance of Alexa 647 dyes. Images were analyzed with ZEN 2012 software with default settings for SIM with minor modifications (Noise filter was set up at -3 to -6 depending on fluorescence intensity). For dSTORM, single molecule blinking was detected using peak mask size of 6-10 pixel, peak intensity to noise set at 5-6 with a x,y 2D Gaussian fit. Overlapping blinking was discarded and X-Y drift was corrected using section based alignment method implemented *in situ*. For 3D imaging, peak mask size was changed to 19-21 pixel, peak intensity to noise set at 5.5 and processed with an experimental PSF determined separately.

**dSTORM and Dual color dSTORM**

**Microscope set-up**

All dSTORM imaging was conducted using a custom-built microscope, based on a fully motorised Olympus IX81 base. Four lasers (100 mW 405 nm Coherent Obis, 100 mW 488 nm Coherent Sapphire, 150 mW 561 nm Coherent Sapphire and a 150 mW 642 nm Toptica iBeam Smart), each with their own shutter control, were expanded to the same diameter and combined using a series of dichroic mirrors into a single free-space beam. Half-wave plates were used to adjust the polarisation before passing the beams through an Acousto-Optical Tunable Filter (AA Optoelectronics) to quickly modulate laser power. The combined beams were again expanded and launched into a single-mode optical fibre (Thorlabs PM-S405-XP) using an Olympus 10× (Air 0.1 N.A.) objective lens. The output of the optical fibre was collimated using an achromatic parabolic mirror collimator (Thorlabs) and passed through a quarter-wave plate to circularly polarise the beam. The free beam then passed through the "TIRF" lens (L1, Thorlabs 200 mm achromatic doublet) and was focused directly onto the back focal plane of the objective lens. This entire subsystem was mounted on a micrometer translation stage to adjust the TIRF angle. The beam was reflected to the objective lens via a multi- edge dichroic filter (Semrock Di01-R405/488/561/635-25×36). The objective lens was an oil-immersion Olympus 1.49 N.A. 100× TIRF apochromatic objective lens. Actively cooled electronmultiplying charged coupled device (EMCCD) cameras (Andor iXon Ultra) were coupled to the camera port of the microscope via an additional 1.5× magnifying relay to achieve optimal Shannon-Nyquist sampling. An additional dichroic mirror (Semrock FF560-FDi01-25×36) in the 4f relay was used to simultaneously image the second colour. Bandpass filters (BP1: Semrock FF01-520/35-25 and BP2: Semrock FF01-670/73-25) in front of the two cameras selected the appropriate dye emission. Typically, camera acquisition was at 33 Hz at full frame (512×512). Laser shutter, acousto-optical tunable filter (AOTF) and camera firing were synchronised using a Data Translation DT9834 data acquisition (DAQ) module, using the internal clock to provide synchronised TTL pulses. Sample positioning was controlled via a motorised micrometer stage (Physik Instrumente) with a XYZ-Nanopositioning stage (Physik Instrumente). A custom built focus-lock mechanism was used to stabilise drift. All software for microscope control was written in C++ and Python. Sub-pixel localization of single molecules, drift correction and localization precision was calculated as described elsewhere (Lowe, Tang et al., 2015). Briefly, our drift correction involved two steps (i) Real-time focus locking (Using TetraSpec beads (0.1 μm, Molecular Probes)) and (ii) post-imaging translational drift correction (Using tracking algorithm that finds fiducial markers at the sample surface plane over time (Lowe et al., 2015)). Cluster analysis was performed as described using Lama (LocAlization Microscopy Analyzer)(Malkusch & Heilemann, 2016).

**Dual color dSTORM**

Two color dSTORM imaging was performed by labelling Miro and several mitochondrial markers. Briefly, HeLa cells were transfected with $^{GFP}$Miro1 or $^{GFP}$Miro2 for (colocalization experiments with MINOS) or $^{GFP}$Miro1/2 and $^{Myc}$Miro1 (for addressing dimer formation) and 24 h post transfection cells were fixed and permeabilized as described earlier. Cells were labelled with mouse anti-GFP and rabbit Mitofilin/c-myc/MTX1/TOM-20 antibodies followed by Alexa 555 and Alexa 647 conjugated secondary antibodies raised against mouse and rabbit respectively. Samples were mounted in 10% Vectashield with 95% Glycerol containing 50 nM TRIS pH 8.0. Imaging was performed in custom built set up described earlier in a two-step method. First dSTORM images were collected using 647 nm laser followed by 561 nm laser with occasional pulse of 405 nm laser. More than 20000 frames were acquired with an exposure time of 30 ms.

**Electron Microscopy and DAB staining**

Cells were fixed with 2% PFA and 1.5% glutaraldehyde in 0.1 M sodium cacodylate, washed with Tris buffer (50 mM Tris/HCl pH 7.6), prior to being incubated with freshly prepared DAB reaction mix (3.5 mM DAB, 0.02% H2O2 in Tris buffer) for 30 minutes in the dark. The reaction was quenched by further washing with Tris buffer followed by further fixation with 1% osmium tetroxide/ 1.5% potassium ferricyanide. Samples were incubated with tannic acid and then dehydrated and embedded in resin, essentially as described in (Nkwe, Pelchen-Matthews et al., 2016). Serial ultrathin sections were collected on formvar coated slot grids, stained with lead citrate and imaged in a 120kV Tecnai Spirit TEM (Thermo Fisher Scientific) coupled to Morada CCD camera (Olympus SIS).

**PLA image analysis**

Images were acquired on a confocal Zeiss 700 confocal microscope with 63x oil objective (1.4 NA). The field of view was selected based on DAPI staining, ensuring each image contains a similar amount of cells and avoiding bias. Images were digitally captured using ZEN software with excitation 405 nm for DAPI, and 555 nm for the PLA red fluorophore (equivalent to Texas Red). The pinhole was set to 1.5 Airy units creating an optical slice of 1.2 um. Image stacks of 4 slices were acquired with voxel dimensions of 0.199 x 0.199 x 0.578 um^3.

Using ImageJ, a suitable threshold was selected for DAPI and PLA signal and kept constant for all data sets. PLA dots and nuclei were then counted using an automated ImageJ script. Staining was performed 3 times. In each experiment, 6-8 images per condition were acquired, and the results averaged. p-values were calculated from unpaired t-test.

**Analysis of signal distribution of mitochondrial markers using micropatterns**

WT and Miro DKO MEF cells were transfected with Tom70(1-70)$^{GFP}$, TRAK1 and $^{myc}$KIF5C. 24 hours after transfection cells were seeded onto “Y” shaped micropatterned coverslips coated with fibronectin (CYTOO) at 15,000 - 20,000 cells / cm$^2$. Cells were allowed to attach to the permissive substrate for 4 hours and then fixed with PFA 4%. Fixed. Cells were immunostained to visualise endogenous expression of an OMM marker (Tom40 - cyan) and an IMM marker (ATP5$\alpha$ - red). Cells with high anterograde transport activity were selected and stacks were imaged using sub-saturation parameters in the confocal microscope. Sholl analysis of mitochondrial signal distribution was performed using a modification of our custom made ImageJ plugin (Lopez-Domenech, Covill-Cooke et al., 2018, Lopez-Domenech, Higgs et al., 2016). Briefly, absolute signal intensity of both mitochondrial markers was measured in the mitochondrial ROI (generated by thresholding the Tom70(1-70)$^{GFP}$ signal) within shells radiating out from the centre of the cells at 1 μm intervals. The intensity profiles generated for each marker and cell were normalized with themselves and a ratio ATP5α signal divided by Tom40 signal was calculated for each shell of the cell. An average value for each shell was then calculated from all the cells in each genotype and plotted as a function of distance from the centre of the cell. 32 cells for each genotype from 3 different experiments were used for the analysis.

.

| Figure | Antibody used | Dilution | Provider | Clone / Cat# |
|---|---|---|---|---|
| 1F | Ms Miro1 | 1:500 | Atlas | AMAb90852 |
| | Ms Miro2 | 1:1000 | NeuroMab | N384/63 |
| | Rb Actin | 1:5000 | Sigma | A2066 |
| | Rb TRAK1 | 1:500 | Atlas | HPA005853 |
| | Rb TRAK2 | 1:500 | Atlas | HPA015827 |
| | Ms β-tubulin | 1:2000 | Sigma | T5293 |
| | Ms ATP5α | 1:2000 | Abcam | ab14748 |
| | Rb Tom20 | 1:1000 | SantaCruz | sc-11415 |
| | Rb Mitofilin | 1:500 | Elabscience | 10395 |
| | Rb Sam50 | 1:1000 | Atlas | HPA034537 |
| | Rb CHCHD3 | 1:1000 | Atlas | HPA042935 |
| | Rb ApooL | 1:500 | Atlas | HPA000612 |
| | Rb IP3R | 1:500 | Abcam | ab5804 |
| | Go GRP75 | 1:500 | SantaCruz | sc-1058 |
| | Ms VDAC1 | 1:1000 | NeuroMab | N152B/23 |
| 3A | Ms GFP | 1:1000 | NeuroMab | N86/38 |
| | Rb Mitofilin | 1:500 | Elabscience | 10395 |
| | Mo Mitofusin1 | 1:1000 | Abcam | ab57602 |
| | Mo Mitofusin2 | 1:1000 | NeuroMab | N153/5 |
| | Rb anti-Sam50 | 1:1000 | Atlas | HPA034537 |
| | Rb Metaxin1 | 1:200 | Atlas | HPA011543 |
| | Rb anti-CHCHD3 | 1:1000 | Atlas | HPA042935 |
| | Rb Tom20 | 1:1000 | SantaCruz | sc-11415 |
| 3B | Rb IgGs | IP: 1μg/ml | Invitrogen | 10500C |
| | Ms Miro2 | IP: 1μg/ml - WB: 1:1000 | NeuroMab | N384/63 |
| | Rb TRAK1 | 1:500 | Atlas | HPA005853 |
| | Rb Mitofilin | 1:500 | Elabscience | 10395 |
| | Rb Sam50 | IP: 1μg/ml - WB: 1:1000 | Atlas | HPA034537 |
| | Rb CHCHD3 | IP: 1μg/ml - WB: 1:1000 | Atlas | HPA042935 |
| 3C | Ms Miro2 | Miro2 1:150 | NeuroMab | N384/63 |
| | Rb Sam50 | Sam50 1:200 | Atlas | HPA034537 |
| | Rb CHCHD3 | CHCHD3 1: 150 | Atlas | HPA042935 |
| 3F | Rb Sam50 | 1:1000 | Atlas | HPA034537 |
| | Rb CHCHD3 | 1:1000 | Atlas | HPA042935 |
| | Rb Miro1 | 1 : 500 | Atlas | HPA010687 |
| | Ms Miro2 | 1:1000 | NeuroMab | N384/63 |
| 3G | Rb Miro1 | 1 : 500 | Atlas | HPA010687 |
| | Ms Miro2 | 1:1000 | NeuroMab | N384/63 |

| | | | | |
|---|---|---|---|---|
| 4A | Ms Myc (9E10) | 1 : 100 | SantaCruz | sc-40 |
| | Ms GFP | 1:100 | NeuroMab | N86/38 |
| 4B | Ms Myc (9E10) | 1 : 100 | SantaCruz | sc-40 |
| | Ms GFP | 1:1000 (purified) | NeuroMab | N86/38 |
| 5A-C | GFP Nanobody-647 | 1: 100 | Custom made | |
| | Ms Miro2 | 1:30 | NeuroMab | N384/63 |
| | Rb Mitofilin | 1:300 | Abcam | ab137057 |
| | Rb Tom20 | 1:500 | SantaCruz | sc-11415 |
| 5F | Rb Miro1 | 1 : 500 | Atlas | HPA010687 |
| | Rb CHCHD3 | 1:1000 | Atlas | HPA042935 |
| 5G | Rb IgGs | IP: 1μg/ml | Invitrogen | 10500C |
| | Rb Mitofilin | 1:1000 | Elabscience | 10395 |
| | Rb Sam50 | IP: 1μg/ml - WB: 1:1000 | Atlas | HPA034537 |
| | Rb CHCHD3 | IP: 1μg/ml - WB: 1:1000 | Atlas | HPA042935 |
| 5H | Rb Sam50 | 1:200 | Atlas | HPA034537 |
| | Ms Mitofilin | 1:200 | Abcam | ab110329 |
| 5I | Rb CHCHD3 | 1: 200 | Atlas | HPA042935 |
| | Ms Mitofilin | 1:200 | Abcam | ab110329 |
| 6A | Rb CHCHD3 | 1: 200 | Atlas | HPA042935 |
| 6C | Rb IgGs | IP: 1μg/ml | Invitrogen | 10500C |
| | Rb TRAK1 | IP: 1μg/ml - WB: 1:500 | Atlas | HPA005853 |
| | Rb TRAK2 | IP: 1μg/ml - WB: 1:500 | Atlas | HPA015827 |
| | Rb Mitofilin | 1:1000 | Elabscience | 10395 |
| | Rb Sam50 | IP: 1μg/ml - WB: 1:1000 | Atlas | HPA034537 |
| | Rb CHCHD3 | IP: 1μg/ml - WB: 1:1000 | Atlas | HPA042935 |
| 6D and E | Rat GFP | 1:1000 | Nakalai | GF090R |
| | Rb CHCHD3 | 1:300 | Atlas | HPA042935 |
| 7A - D | Rat GFP | 1:1000 | Nakalai | GF090R |
| | Rb anti-Tom40 | 1:500 | Proteintech | 18409-1-AP |
| | Ms anti-ATP5α | 1:500 | Abcam | ab14748 |

**Supplementary Table 1**

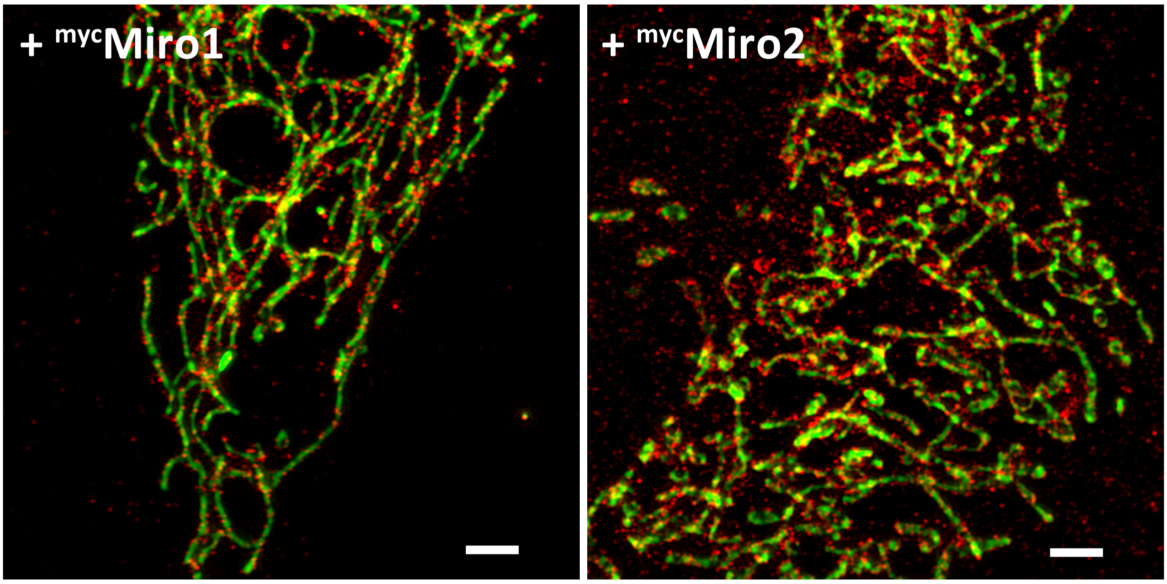

### Supplementary Figure 1

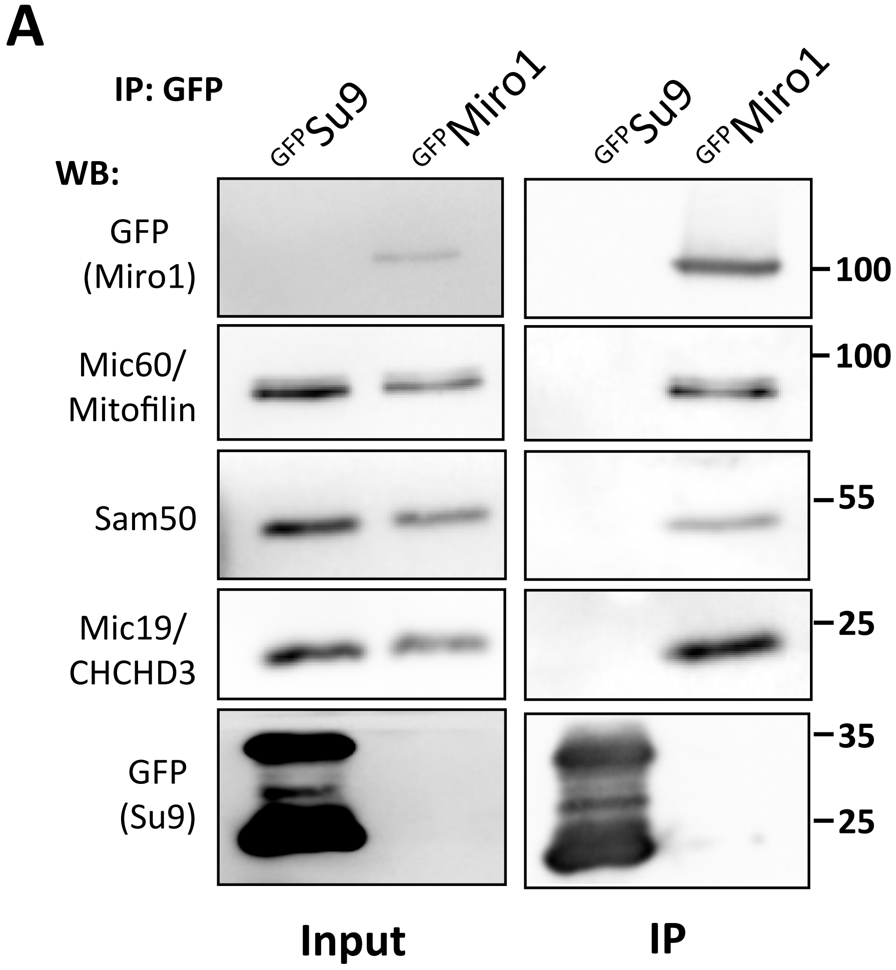

**Supplementary Figure 2**

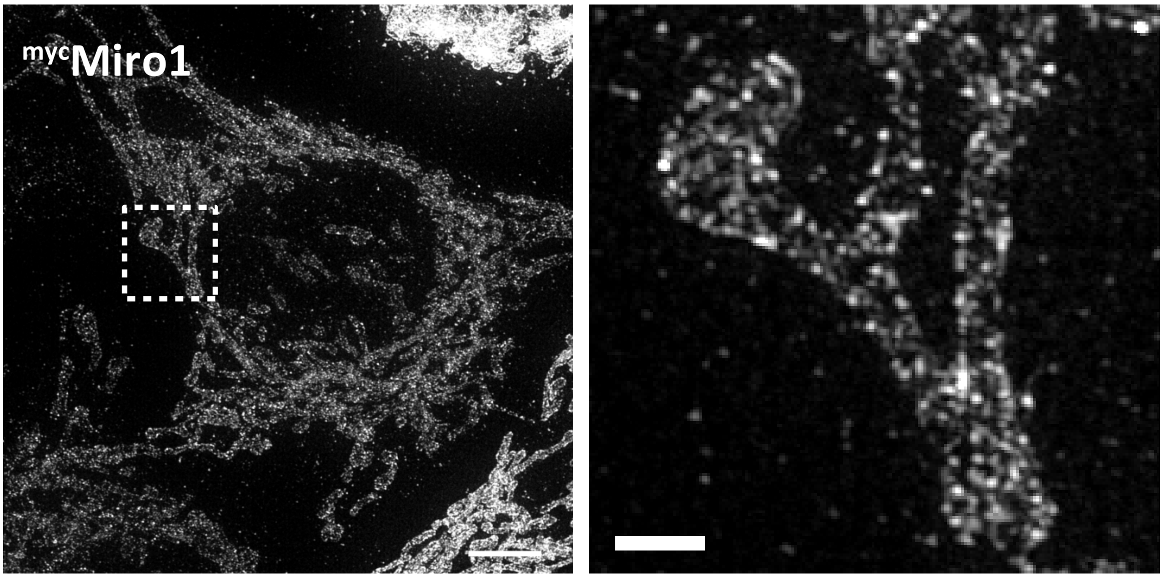

**Supplementary Figure 3**

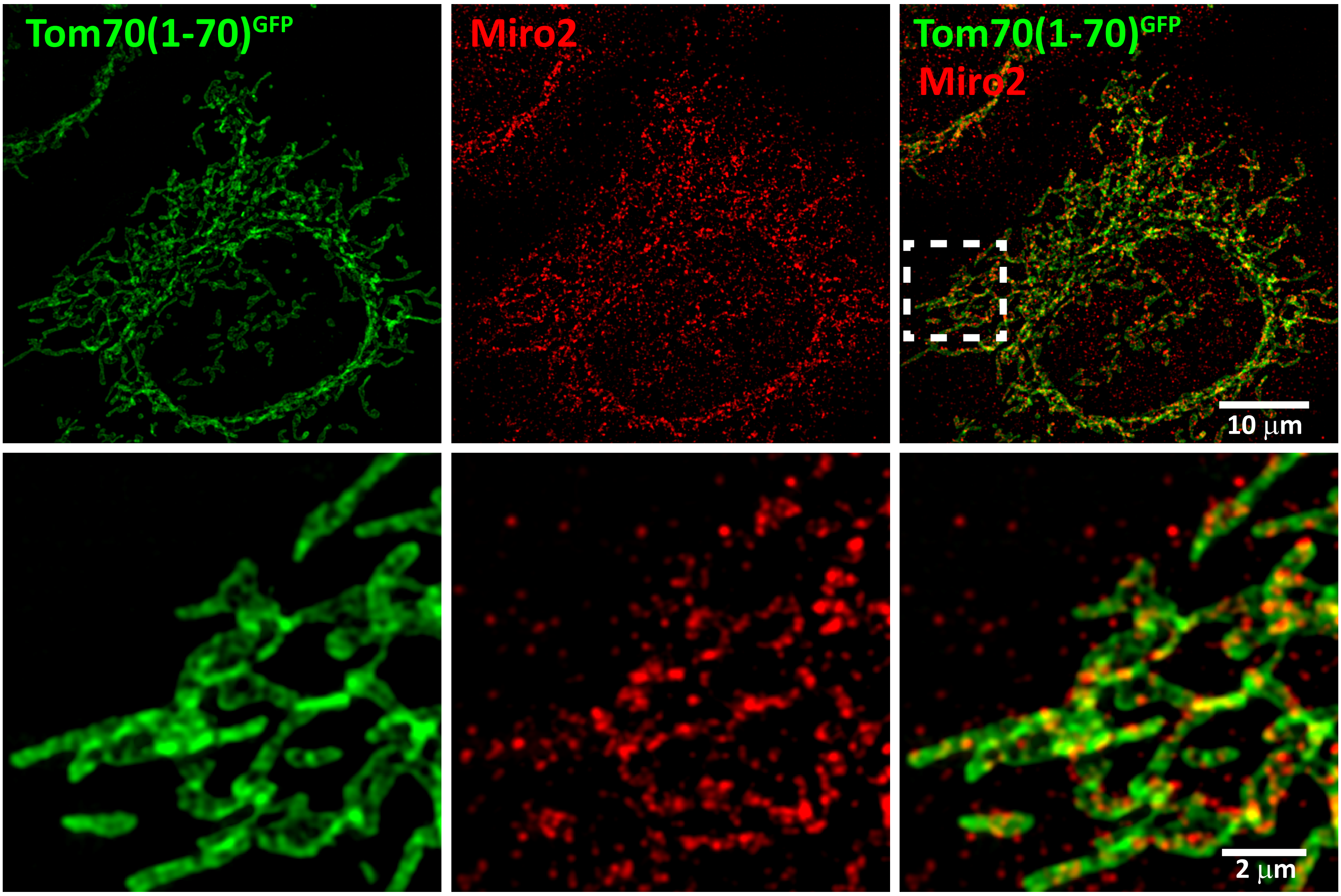

**Supplementary Figure 4**

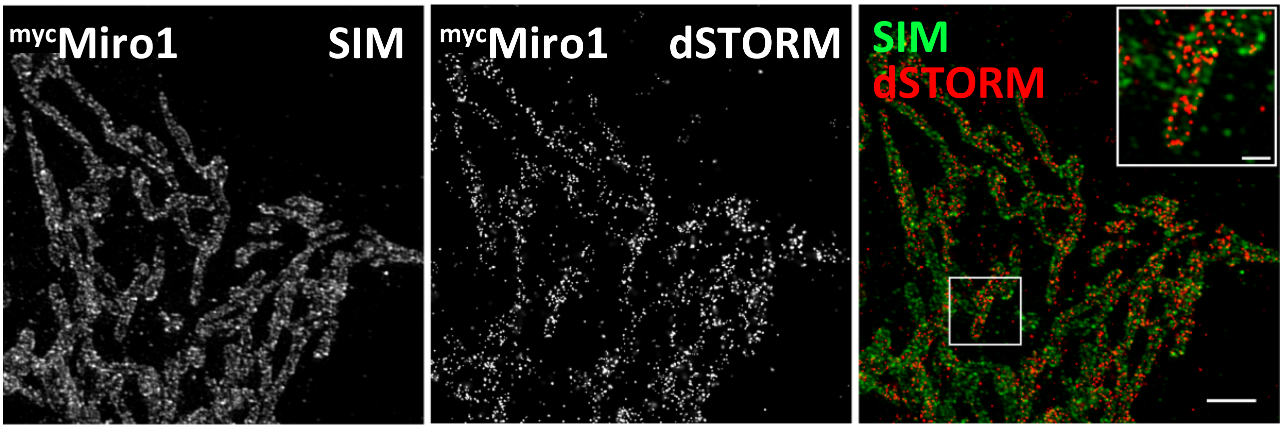

**Supplementary Figure 5**

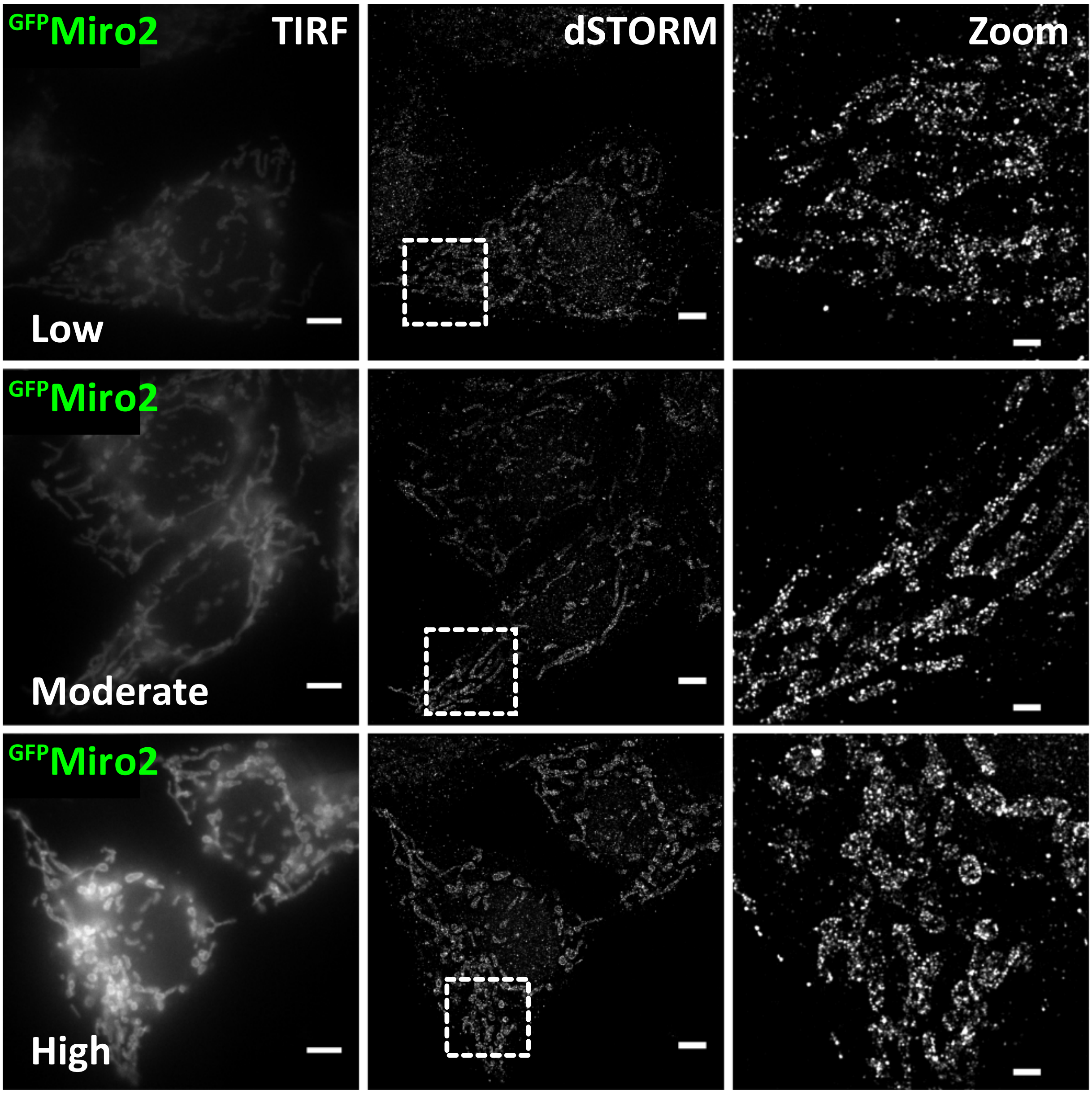

**Supplementary Figure 6**

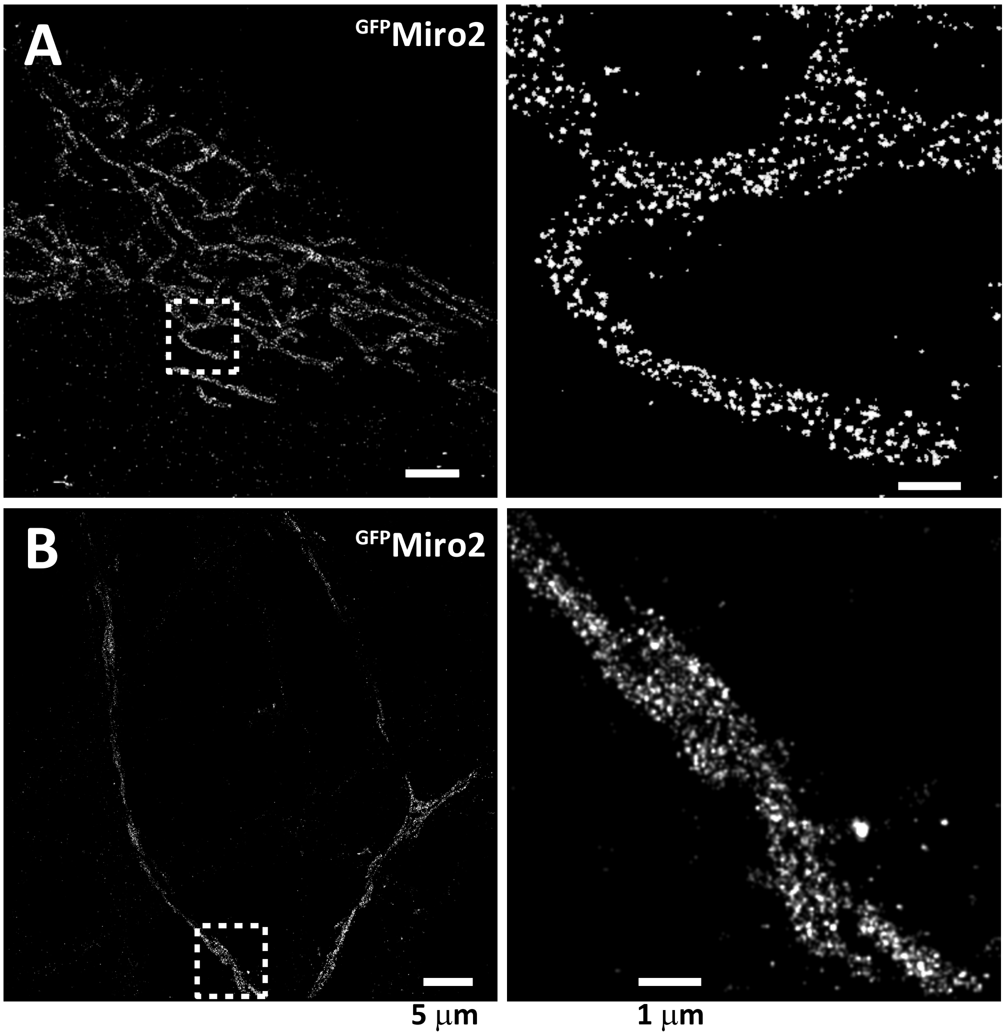

**Supplementary Figure 7**

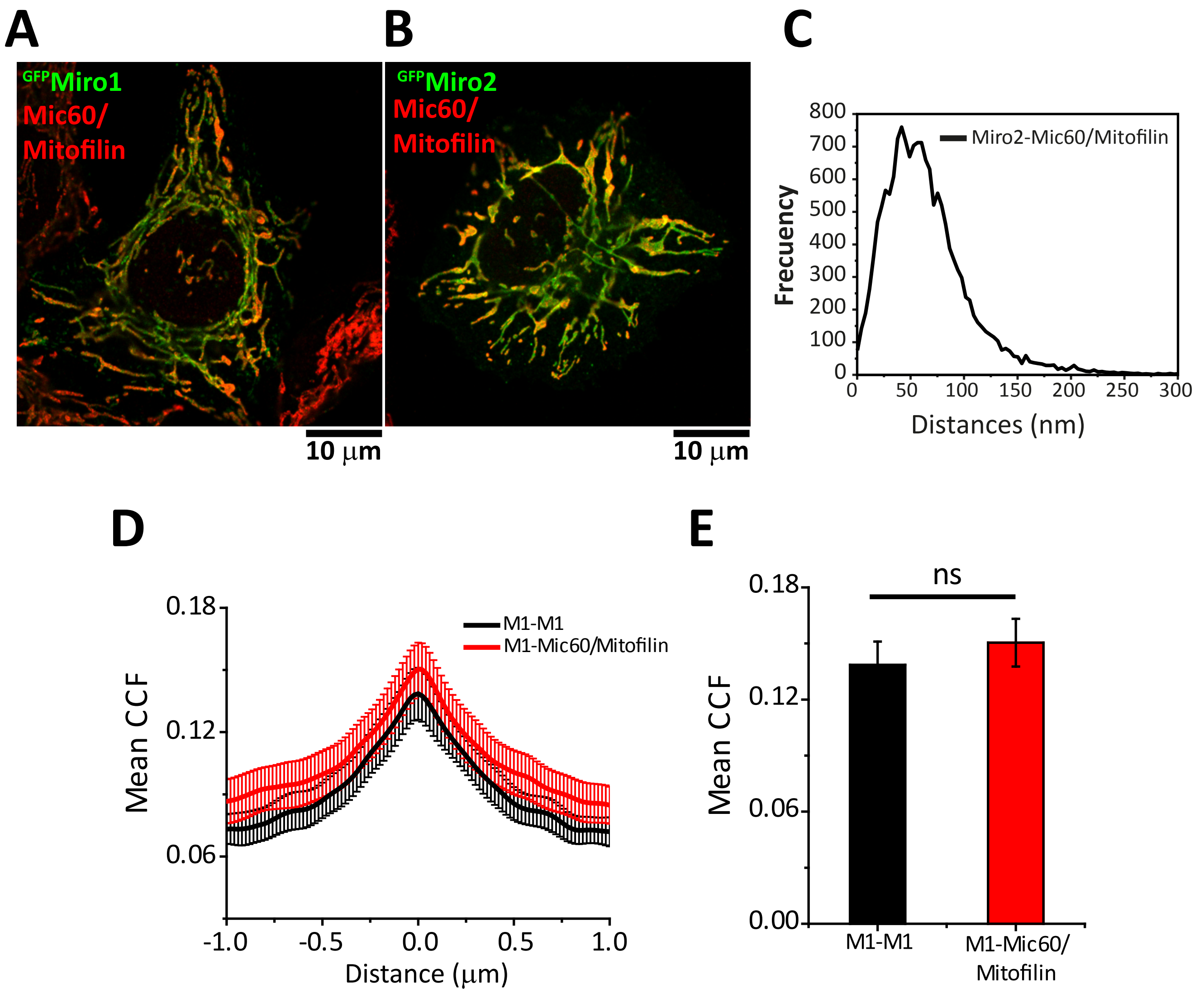

**Supplementary Figure 8**
